# Probing the effect of glycosaminoglycan depletion on integrin interactions with collagen I fibrils in the native ECM environment

**DOI:** 10.1101/2020.02.07.935932

**Authors:** Jonathan Roth, Cody L. Hoop, Jonathan K. Williams, Robert Hayes, Jean Baum

## Abstract

Fibrillar collagen–integrin interactions in the extracellular matrix (ECM) regulate a multitude of cellular processes and cell signalling. Collagen I fibrils serve as the molecular scaffolding for connective tissues throughout the human body and are the most abundant protein building blocks in the ECM. The ECM environment is diverse, made up of several ECM proteins, enzymes, and proteoglycans. The contents of the ECM environment are modulated by disease and aging and may influence these critical collagen–integrin interactions. In particular, glycosaminoglycans (GAGs), anionic polysaccharides that decorate proteoglycans, become depleted in the ECM with natural aging and their mis-regulation has been linked to cancers and other diseases. The impact of GAG concentration in the ECM environment on collagen interactions is not well understood. Here, we integrate protein adhesion assays with liquid high resolution atomic force microscopy (AFM) to assess the affects of GAG depletion on the interaction of collagen I fibrils with the integrin α_2_I domain. Adhesion assays demonstrate that α_2_I preferentially binds to GAG-depleted collagen I fibrils. By amplitude modulated AFM in air and in solution, we find that GAG-depleted collagen I fibrils retain structural features of the native fibrils, including their characteristic D-banding pattern, a key structural motif. AFM fast force mapping in solution shows that GAG depletion reduces the stiffness of individual fibrils, lowering the indentation modulus by half compared to native fibrils. Together these results shed new light on how GAGs influence collagen I fibril– integrin interactions and may aid in strategies to treat diseases that result from GAG misregulation.

**Statement for broader audience:** Aging and disease result in mis-regulation of glycosaminoglycan (GAG) levels in the extracellular matrix (ECM), which may affect fibrillar collagen interactions that are vital for cellular processes. Here, we characterize the impact of GAG depletion on collagen–integrin α_2_I domain interactions and collagen fibril topography and stiffness. We show that GAG depletion increases collagen–α_2_I binding and reduces stiffness in comparison to native fibrils. These results may inform on strategies for treating GAG mis-regulation.

## Introduction

Intermolecular interactions between fibrillar collagens and collagen-binding integrins in the extracellular matrix (ECM) drive numerous cellular processes, including cell adhesion and motility, platelet aggregation, cell development, differentiation, and hemostasis (1–3). While much knowledge has been reaped from studying integrin binding to triple helical collagen model peptides, the ECM is quite complex, composed of collagen fibril assemblies and also other ECM proteins, matrix metalloproteinases, and proteoglycans (PGs), which contain glycosaminoglycan (GAG) polysaccharides (4) (Figure 1A). As part of the natural aging process, GAGs are depleted in the ECM (5,6). How this GAG depletion alters the functional activity of collagen fibrils is not well understood. In this study, we aim to understand how GAG depletion impacts the critical interactions between collagen fibrils and integrins.

**Figure 1.**
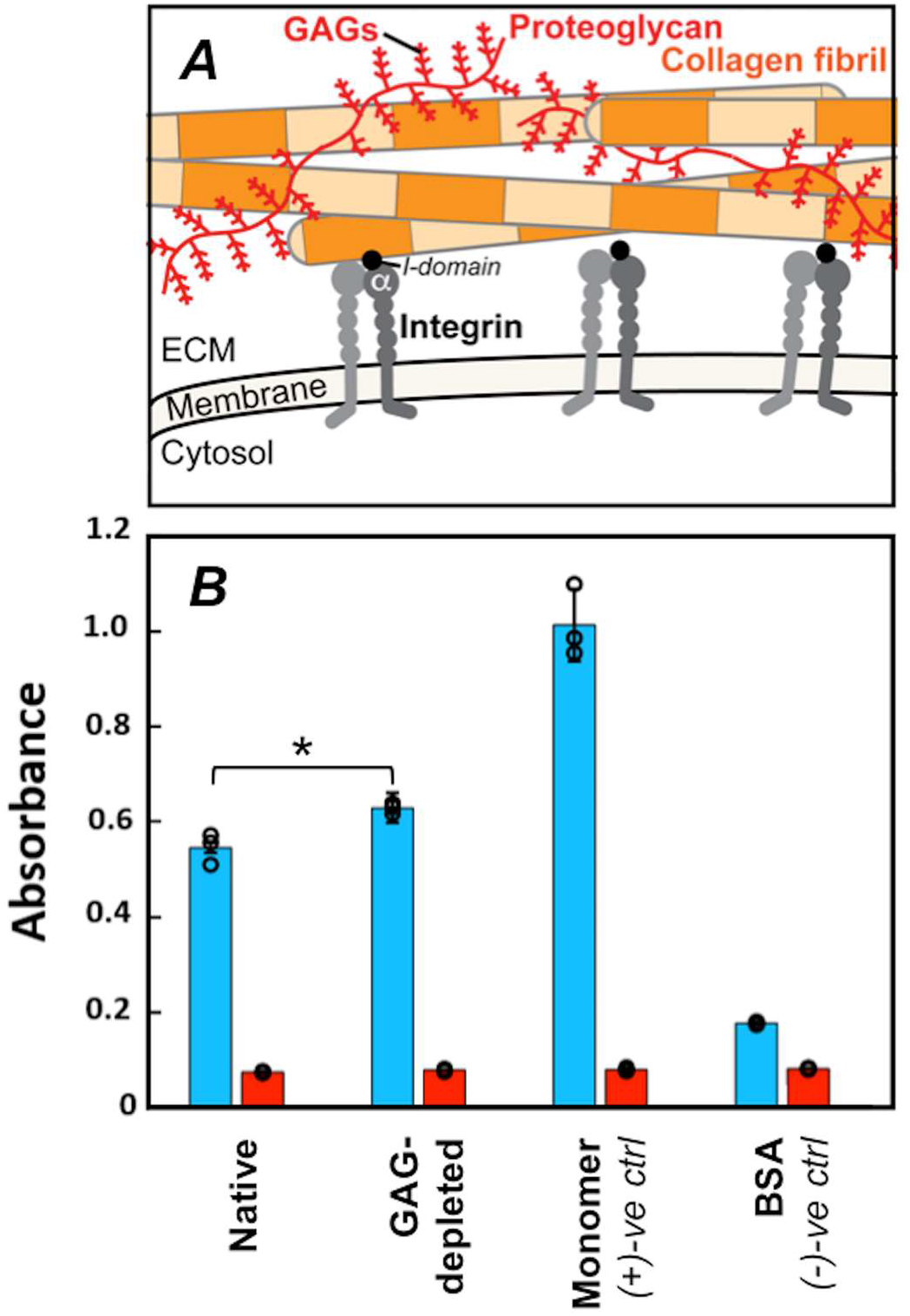
Collagen I fibril–integrin α_2_I interactions in the native ECM environment. (A) Cartoon highlighting multiple components of the ECM. In this study, we focus on collagen I fibrils (orange); integrin cell surface receptors (gray), specifically the I-domain (black) of the α-subunit (αI); and glycosaminoglycans (GAGs), which are polysaccharide chains displayed on proteoglycans (red). Integrin αI domains and GAGs interact with individual collagen I fibrils in the ECM. (B) ELISA adhesion assay of integrin α_2_I binding to collagen I in varying contexts (native: collagen I fibrils from rat tail tendon ECM in PBS, GAG-depleted: collagen I fibrils from ECM with 85% GAG depletion in PBS, monomer: purified rat tail tendon collagen I in 10 mM acetic acid, and BSA: bovine-serum albumin in PBS.) A one-way ANOVA analysis confirmed that there is a significant difference between α_2_I binding to native and GAG-depleted collagen I fibrils with a p value of < 0.05. Each binding condition was performed in the presence of either 5 mM MgCl2 (blue) or 5 mM EDTA (red). Error bars indicate the standard deviation of the measurements, which were taken in triplicate. Individual measurements are shown as circles.

The most abundant fibrillar collagen in the human body is collagen I, primarily found in tendons, bone, and skin (7,8). Collagen I self-assembles into fibrillar structures that are up to hundreds of nanometers in diameter and hundreds of micrometers long (9–11). These fibrils are made up of triple helical monomer units, heterotrimers of three protomer chains (9–11). The uniform staggered arrangement of triple helical monomers within the fibrils gives the fibrils a banded topography defined by regions of low and high protein density, referred to as the gap and overlap regions (9–11). This repeating pattern, called D-banding, has a characteristic periodicity of 67 nm that can be observed by high-resolution microscopy methods (9–11). These fibril units serve as the main building blocks of collagen in the ECM and act as the molecular scaffolding for cells that make up connective tissues (9–11). Collagen I is highly biologically active, interacting with numerous ECM biomolecules (Figure 1A) to enable cell signaling, regulate collagen turnover, and facilitate many cellular processes (7,8,12,13).

Integrins are an important class of collagen interaction partners for cell signaling (1). One of the most widely distributed collagen-binding integrins, integrin α_2_β_1_ is primarily found on platelets and epithelial cells (1–3,14). Integrin α_2_β_1_ interacts with collagen I via a divalent metal coordination of its metal ion-dependent adhesion site (MIDAS) on the inserted (I) domain of the α_2_ subunit (α_2_I) with a glutamate in a GXX’GEX” motif in collagen I, where X’ is often hydroxyproline (O) and X” is R or N (15–18). The α_2_I domain retains its structure and interaction affinity for collagen I in isolation (16). Thus, it can be used as a biologically active mimic for collagen I–integrin α_2_β_1_ interactions. Previously, our lab and others have studied collagen–integrin αI interactions in the context of triple helical collagen model peptides (CMPs) (16–26). Farndale and co-workers have developed a library of CMPs to determine the specificity of integrin αI domains for collagen binding motifs (27). The GFOGER sequence was identified as a high affinity motif in collagen I (17), and a number of lower affinity sites were identified with a canonical sequence of GXOGER/N (15,25). We have investigated how collagen–αI interactions are perturbed upon modulation of these peptide sequences or in the presence of disease-affiliated substitutions (19–22). However, in the ECM, collagen I triple helical monomers assemble into fibrils. In these CMPs and collagen monomers, all possible integrin binding sites are fully exposed. Yet, upon assembling into fibrils, many binding sites become buried within the fibril core and seemingly inaccessible from the fibril surface. We have previously proposed that molecular motions on the collagen I fibril surface may facilitate increased surface accessibility to some integrin recognition sites that are otherwise hidden by the complex fibril architecture (28).

In the ECM, the highly conserved collagen assembly is regulated by PGs, in which protein cores are bound to anionic polysaccharide chains, called GAGs (29,30). A large diversity of PGs and GAGs are distributed across connective tissues for proper ECM activity (29,30). In all tissues, PGs recognize a number of specific amino acid sequence motifs on collagen fibrils (29) to act as part of the ECM scaffolding. In addition to their structural roles, GAGs have been shown to be essential for wound healing, cell adhesion, and cell motility (31–34). As humans age, the concentration of GAGs in the ECM becomes depleted (29,35,36). However how changes in the ECM environment impact critical fibrillar collagen I-α_2_I interactions has not been elucidated. Here, we probe how alterations in the GAG content in the ECM modulate integrin α_2_I interactions with native collagen I fibrils in solution.

In order to interrogate these critical functions in a physiological environment, we investigate fibrillar collagen I-integrin α_2_I interactions in the native ECM environment. Given that collagenous tissues function in a fully hydrated environment, it is important to investigate α_2_I binding to the fibrillar form of collagen in solution. In this study, we probe the influence of GAG concentration on the 1) collagen-integrin α_2_I binding activity using an enzyme-linked immunosorbent assay (ELISA) and on the 2) structure and mechanics of collagen I fibrils through advanced AFM techniques in solution. Together, these results provide a better understanding of how GAG depletion, an inevitable result of aging, impacts the interactions of collagen I fibrils with its cell receptor, integrin α_2_I, in its native environment.

## Results

### Assessing the impact of GAG depletion on collagen–integrin α_2_I domain interactions

The complexity of the ECM makes it critically important to investigate collagen I fibrils in the context of their native environment. Rat tail tendon provides a rich source of collagen I fibrils in an intact ECM that also includes other ECM proteins, proteoglycans, and GAGs. The environmental contents of the ECM may influence the functional activity of the collagen I fibril scaffold within it. Interactions of collagen I fibrils with the integrin α_2_I domain is critical for platelet adhesion. To determine the impact of GAG depletion on collagen-α_2_I interactions, we performed ELISA adhesion assays of recombinant α_2_I to native collagen I fibrils, GAG-depleted (85% GAG reduction) collagen I fibrils, and positive and negative controls. As the collagen I-α_2_I interaction is dependent on coordination of a divalent metal cation, the assay is performed in the presence of Mg^2+^ or ethylenediaminetetraacetic acid (EDTA). Minimal α_2_I adhesion is observed in the presence of EDTA (Figure 1B); therefore all binding events are metal dependent, as expected. Bovine serum albumin (BSA) was used as a negative control, as no specific α_2_I interaction is expected. Acid solubilized, purified collagen I triple helical monomers serves as a positive control for collagen I-α_2_I binding. We see a high level of α_2_I binding to this triple helical collagen I monomer, consistent with the view that all integrin binding sites are fully exposed. For the native collagen fibrils, the integrin α_2_I domain shows less adhesion relative to collagen triple helical monomers (Figure 1B). This has been shown previously by other labs, which attribute this reduced adhesion to the reduced accessibility of integrin binding sites (37), many of which are bundled into the fibril core upon fibril assembly. GAG-depleted fibrils, relative to native fibrils, show slightly higher binding to integrin α_2_I (Figure 1B). The adhesion of α_2_I to GAG-depleted fibrils is trending toward the fully exposed triple helical monomer. This may suggest that integrin binding sites within the GAG-depleted collagen I fibrils are becoming more accessible upon removal of GAGs. Thus, the ability of integrin α_2_I to interact with collagen I fibrils is modulated by GAGs in the ECM environment, and the regulation of GAG concentration is imperative for proper collagen activity. In order to better understand how the presence of GAGs modulates collagen I fibril–α_2_I interactions, we probed how GAG content modulates the structure and mechanics of the individual collagen I fibrils.

### Characterizing native collagen I fibrils in solution

The emergence of sophisticated AFM imaging modes has enabled direct, high-resolution images of structure or mechanical/surface properties of biological systems *in situ* under physiological aqueous conditions (38,39) We characterized the topography of intact collagen I fibrils extracted from rat tail tendon in air and in solution by amplitude-modulation (AM-) AFM. Collagen I fibrils are expected to display a canonical repeating 67-nm banding pattern that is produced by the staggered arrangement of triple helical units, referred to as D-banding (40). AM-AFM confirms such banding is present in rat tail tendon collagen I fibrils adsorbed to a glass slide in air and when immersed in PBS buffer solution (Figure 2). Here, the sinusoidal periodicity in the height along the main fibril axis is observed in both imaging environments, with alternating high (“overlap,” yellow), and low (“gap,” red) regions. Height profiles extracted from the apex of the collagen I fibrils in air reveal a distinct sinusoidal oscillation of overlap and gap regions (Figure S 1A,C). Although this periodicity in the height profiles is not as clear from images in PBS, it is discernible in the phase channel (Figure S2A–B). Two-dimensional Fourier transforms of the phase channel from imaging native collagen I fibrils revealed an average periodicity of 65.6 ±1.0 nm (N=5) in air and 65.8 ± 0.8 nm (N=5) in PBS (Table I). Thus, the fibril periodicity, a structural hallmark of fibrillar collagens, is maintained in an aqueous environment.

**Figure 2.**
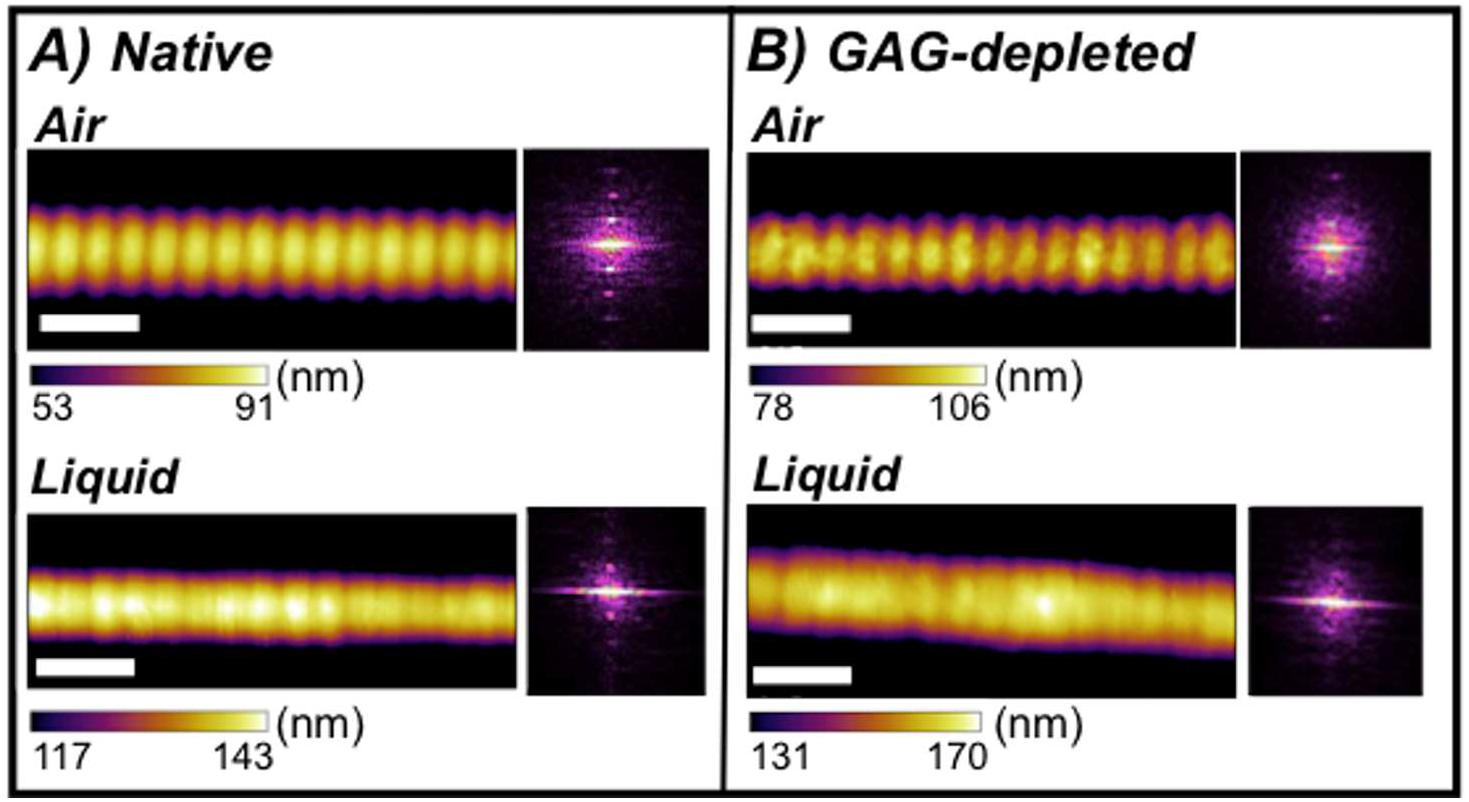
AM-AFM imaging of collagen I fibrils. Collagen I fibrils bear a regular, characteristic D-banding pattern that is produced by the staggered arrangement of triple helical collagen monomers in the fibril. This D-banding is resolved here by AM-AFM experiments for both (A) native and (B) GAG-depleted collagen I fibrils imaged in air and liquid environments. The white scale bars are 200 nm in each image. The Fourier transform insets show the periodicity of each fibril and scale bar ranges were chosen to accentuate the D-banding in the images. Uncropped height, amplitude, and phase images are shown in the SI (Figure S2, S3).

**Table I:** Overview of results for the control and chondroitinase treated fibrils. Visualization of the distribution of these values are shown in the SI (Figure S7).

|  | <u>**Native**</u> | <u>**GAG-depleted**</u> |
| --- | --- | --- |
| **Periodicity (air)** | 65.6 ± 1.0 nm | 66.0 ± 0.2 nm |
| **Periodicity (liquid)** | 65.8 ± 0.8 nm | 66.2 ± 1.0 nm |
| **Overlap-Gap**<br>**Height Difference (air)** | 3.6 ± 2.4 nm | 3.6 ± 2.7 nm |
| **Hydration Ratio** | 1.79 ± 0.06 | 1.9 ± 0.2 |
| **Modulus (liquid)** | 6.4 ± 1.4 MPa | 2.9 ± 1.0 MPa |

### The fibril nanoscale structure is maintained upon depletion of GAGs

We then asked if this characteristic structure of individual collagen I fibrils is impacted by the concentration of GAGs in their environment. We used AM-AFM to probe perturbations to the topography of collagen I fibrils in ECM with a 76% reduced GAG concentration upon treatment with chondroitinase. GAG-depleted collagen I fibrils display D-band periodicities of 66.0 ± 0.2 nm in air (N=5) and 66.2 ± 1.1 nm in PBS (N=4), as measured from 2D Fourier transforms of the phase channel (Table I). This is consistent with the native fibril data and with previously reported AFM results of chondroitinase-treated tendons in air (41).

Another assessment of fibril surface topography is the overlap–gap step height. This is the difference between the maximum height of the overlap and the minimum height of the gap (Figure S4). Upon GAG depletion, the overlap–gap step height also remained unchanged, as chondroitinase-treated collagen I fibrils exhibit an overlap–gap step height of 3.6 ± 2.7 nm in air compared to 3.6 ± 2.4 nm in native collagen I fibrils (Figure S7A). Step-heights in liquid were unable to be determined as gap and overlap regions were indistinguishable across height profiles (Figure S1B, D). Overall, the data show that GAG association does not significantly impact fibril periodicity or topography. This indicates that the collagen I fibril nanoscale surface structure is preserved upon GAG depletion in aqueous solution.

### AFM force mapping reveals a reduction in stiffness of individual collagen I fibrils upon GAG depletion relative to native fibrils

Although GAG depletion does not significantly affect the surface structure of collagen I fibrils, we asked if the mechanical properties of the fibrils were perturbed upon GAG depletion. It has previously been proposed that mis-regulation of GAGs perturbs the mechanics of the ECM, leading to a loss of biological function and aberrant cell activity (42). The mechanical properties of collagen impact its molecular interactions, and collagen stiffness has been shown to direct cell movement and determine cell fate (43,44). However, the mechanism and extent to which GAGs alter the mechanics of individual collagen I fibrils has not been elucidated. We used fast force mapping (FFM-) AFM to determine the indentation modulus, a measure of local nanoscale stiffness, of individual collagen I fibrils in solution in native and GAG-depleted states. Figure 3 presents a 1 x 1 μm force map of native and GAG-depleted collagen I fibrils, respectively. Unlike in Figure 2, which is an amplitude-modulated height image, it is important to emphasize that these images are 2D maps displaying the indentation modulus of each pixel (Figure S6). This enables mechanical data to be assessed at different locations within the D-bands along the fibrils.

**Figure 3.**
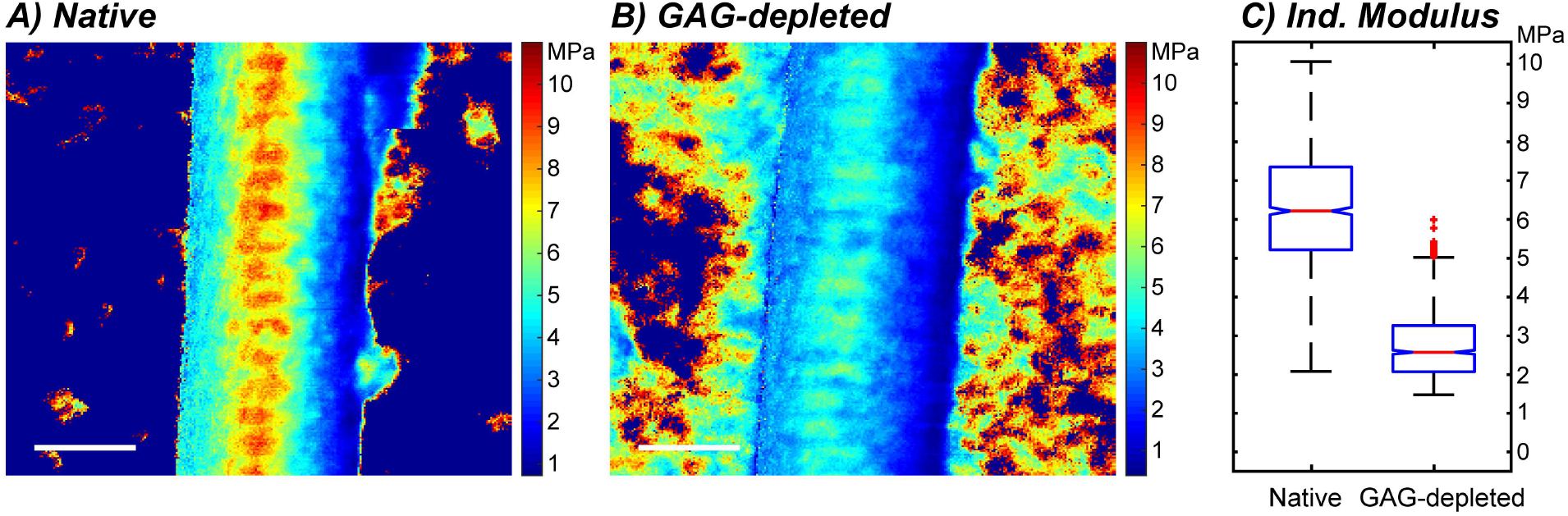
Nanoscale indentation modulus maps of collagen I fibrils in solution. (A,B) Representative 1 x 1 μm fast force maps of individual (A) native and (B) GAG-depleted collagen fibrils from FFM-AFM experiments. The color at each pixel shows the *local* nanoscale indentation moduli (1 pixel / every ~15 nm^2^). The FFM images are spatial reconstructions of 65,536 discrete force–distance curves that were fitted using standard indentation protocols (57) and subsequently replotted in Cartesian space using in-house MATLAB code. A large scan size was selected to capture more than ten D-bands per image, enabling meaningful comparisons between internal regions, fibrils, and GAG concentration. White scale bars indicate 200 nm. (C) Box and whisker plots compare a subset of indentation moduli data in (A) and (B) for force curves acquired only at the apex down the fibril long axes. In (C), data for five fibrils in each condition are presented for increased statistical confidence. A one-way ANOVA analysis confirmed that there is a significant difference between the conditions with a p value of < 0.01.

Notably, D-banding is evident in both force maps; the overlap region systematically has a larger indentation modulus than the gap regions, leading to a distinct light/dark/light patterning every ~67 nm down the long axis. While the demarcation between gap/overlap is less well-defined than for AM-AFM data, the results show that even in solution, collagen mechanics oscillate as dictated by the underlying D-banded structure. Specifically, the staggered arrangement of triple helical monomer units leads to a more elastic gap region compared to overlap regions due to the reduced molecular density. Previous studies have reported a 20% increase in modulus of the overlap relative to the gap as measured by fast nanoindentation and attributed it to the 20% higher protein density in the overlap relative to the gap regions (45).

Native and GAG-depleted collagen I fibrils display important differences in the forces measured. A significant reduction in average local indentation modulus is evident upon GAG depletion relative to the native fibril, while preserving the native D-banded structure. In Figure 3A, overlap regions of the native fibril have moduli of ~9 MPa *(red)* compared to ~5 MPa *(light blue)* for GAG-depleted fibrils. Likewise, corresponding gap regions shift from ~7 MPa (yellow) to ~2 MPa *(dark blue)* upon GAG depletion (Figure 3B). To better understand these differences we focused on the 256 force curves along the apex of each fibril (Figure 3C). The native collagen I fibrils exhibit a mean indentation modulus of 6.4 ±1.4 MPa with a broad distribution of values over the fibrils measured, using an indentation speed of 16 μm/second (Figure 3A, C, S7C). This is consistent with reported indentation moduli of collagen I fibrils with similar indentation speeds in solution (45). For GAG-depleted fibrils, both the average indentation modulus (2.9 ± 1.0 MPa) and distribution is reduced (Figure 3B, C), indicating that GAG depletion reduces fibril stiffness relative to the native fibrils.

## Discussion

With an increasingly aging population, it is important to understand the impact of GAG depletion on interaction processes in the ECM of connective tissues. Not only is GAG depletion significant in aging, but mis-regulation of GAGs in the ECM has been linked to various cancers, diabetes, fibrosis, and other diseases (46–50). Thus, it is important to understand how the concentrations of GAGs in the ECM influence molecular interactions that drive cellular processes. Here we address how the critical interactions between individual collagen I fibrils and the integrin α_2_I domain are impacted by a reduction in GAGs in the native ECM. We show that GAG depletion reduces the indentation modulus by a factor of two-to-three relative to the native fibril and that the reduction is apparent across the length of the entire fibril. The breadth of the reduction is consistent with the identification of PG binding sites across broad regions of both the gap and overlap regions within the fibril structure (31–34,51).

Collagen–integrin interactions play a critical role in numerous cellular processes such as cell development and differentiation, hemostasis, and platelet aggregation and have been the subject of numerous studies involving disease, such as cancer (1–3,14,52,53). Collagen binding integrins have been shown to bind directly with individual collagen fibrils through the α_2_I domain (37). Extensive studies using triple helical model peptides have identified unique binding sites to α_2_I (15,17,18,24). These binding sites are fully exposed in triple helical peptides or the full-length collagen I monomer. However, when the monomers become incorporated into the fibril, many of these integrin binding sites become buried within the fibril core. Here, we investigate collagen–integrin interactions in their native context of the ECM to understand how environmental factors influence these interactions.

Our data, using ELISA and liquid high-resolution AFM, indicate that GAG depletion within the ECM impacts on both the stiffness of the collagen I fibrils as well as the binding to the α_2_I domain of integrin relative to the native fibril. We propose that enhanced integrin binding to fibrillar collagen in response to reduced GAG concentration may arise from two different factors or a combination thereof. One proposal is that diminished GAG concentration on the surface of the collagen fibrils results in increased accessibility of α_2_I to collagen binding sites as a result of increased fibril exposure. It has been shown via detailed mapping of interaction sites onto the fibril structure that some PG binding regions overlap with integrin binding sites (32). Therefore, it is possible that the interaction of GAGs with the collagen I fibril may interfere or compete with binding by integrins. A second proposal is that the 2-3 fold reduction in indentation modulus and reduced stiffness of the GAG-depleted collagen I fibrils relative to the native state may modify the inherent binding affinity of α_2_I to collagen. One aspect by which inherent α_2_I binding may be modulated is that reduced indentation may promote a higher propensity for deformation within the collagen fibril. Despite our observation that nanoscale topography of collagen I fibrils does not change upon GAG-depletion, this deformation propensity may allow rearrangement of monomers within the fibril and grant α_2_I access to binding sites within the core that are otherwise inaccessible. Integrin binding sites that are exposed in the collagen model peptides are proposed to be buried and inaccessible from the fibril surface in the complex collagen fibril architecture (37,54). We have previously proposed, through molecular dynamics simulations on the nanosecond timescale that molecular rearrangements may facilitate access to otherwise hidden binding sites of integrin (55) and that monomer fluctuations within the fibril may play a role in regulating collagen-integrin interactions.

In conclusion, we have shown that under physiological solution conditions, GAG depletion enhances collagen-integrin α_2_I domain interactions while decreasing the indentation modulus of individual collagen I fibrils relative to native fibrils and maintaining their native nanoscale topography. It is increasingly clear that alteration of GAG concentration is intimately associated with the progression of aging (5,6), which leads to disruptions in the functional landscape of the ECM. Still, further research will be required to make a direct association between GAG depletion and cell activity or how the effect propagates to longer length scales beyond the fibril building block. The current study provides new knowledge of how GAG depletion affects collagen I fibril-integrin interactions and how it modulates collagen I fibril structure and mechanics. This expands on the understanding of how aging processes, namely alterations in GAG levels, manifest in the ECM and may aid in the development of new treatment strategies.

## Materials and Methods

### Chondroitinase treatment of rat tail tendons

Rat tail tendons were harvested from frozen–thawed rat tails. Chondroitinase treatment of tendons was performed in a similar manner to previously published methods (41). In short, sections of tendon several centimeters in length were cut from the exposed tail tendons. For chondroitinase treatment, the tail tendon sections were placed into a 4 mL solution of 0.15 U/mL chondroitinase ABC (Sigma-Aldrich) in 0.1 M sodium acetate and 0.1 M Tris-HCl, pH 8. For the untreated control, tendons were placed in the same buffer in the absence of chondroitinase. These are referred to as native fibrils. The tendons were incubated in their respective solutions at 37°C overnight. After incubation, then tendon chunks would then be treated for AFM imaging, GAG quantification, or the ELISA assay.

### Quantification of GAG depletion

GAG depletion was quantified by using the dimethylmethylene-blue (DMMB) assay (56). Tendons, prepared in triplicate, were then rinsed with water and placed into 1 mL of a papain solution (500 μg papain, 0.1 M dibasic sodium phosphate, 0.01 M EDTA, 0.0144 M L-cysteine) at 60°C for 24 hours to digest the tendons. The resulting solution was stained with DMMB and absorbance was measured at 525 nm. A standard curve of varying chondroitin sulfate concentrations (0.25–1.5 μg/mL) was used to quantify the GAG concentration in the treated and control samples.

### Integrin α_2_I domain expression and purification

The integrin α_2_I domain (residues 142-336) was recombinantly expressed in *Escherichia coli* BL21(DE3) cells with an N-terminal His6 tag by induction with 1 mM of IPTG for 16 hours at 25° C to stimulate protein production and then harvested. The cells were lysed with a 20% sucrose TES buffer. The His6-α_2_I domain was purified using a Ni^2+^-charged HisTrap HP (GE Healthcare Life) and buffer exchanged to PBS, pH 7.4 with a PD-10 desalting column (GE Healthcare Life). Protein concentration was determined by measuring A280 with a molar extinction coefficient of 20,400 M^-1^cm^-1^.

### ELISA Assay

Following incubation, native and GAG-depleted tendon sections of similar weight (±0.1 mg) were placed into 1.5 mL of PBS and subjected to five minutes of tearing by tweezers and sonicated three times for 10 seconds at a 50% amplitude. The tendons were then manually pulled apart again by tweezers for two and a half minutes and sonicated for 15 seconds at 50% amplitude to create a solution of dispersed fibrils.

Immulon 2HB 96-well plates (Thermo Scienfic) were coated with triplicate samples of native and GAG-depleted rat tail tendon collagen dispersions as prepared above and purified collagen I monomer (Corning, 10 μg/mL in 20 mM acetic acid, positive control) overnight at 4° C. The following day the solutions that did not adhere to the microplates were discarded. The wells were then blocked for 1 hour with 200 μL of 50 mg/mL BSA in PBS at room temperature. Three blank wells were also coated with 200 μL of 50 mg/mL BSA in PBS for 1 hour at room temperature as a negative control. The wells were then washed three times with a solution of PBS with 200 μL of 1 mg/mL BSA containing either 5 mM MgCl2 or 5 mM EDTA. Recombinant integrin α_2_I domain was then added to bind to the plated collagens or BSA for 1 hour at room temperature. After washing three times with the MgCl2 or EDTA buffer, 100 μL of mouse anti-integrin α_2_I antibody was added to the wells at a 1:2000 dilution for 45 minutes followed by binding 100 μL of a goat HRP-conjugated anti-mouse antibody at a 1:5000 dilution for 30 minutes. Washing steps were performed between each binding addition. A TMB Substrate Kit (Pierce) was then used according to the manufacturer’s protocol to measure integrin α_2_I binding activity. Absorbance was measured at 450 nm.

### Atomic force microscopy (AFM)

For AFM sample preparation, following incubation, the tendon sections were rinsed with fresh PBS buffer solution (pH 7.4) and smeared onto a microscope glass slide in order to deposit collagen fibrils onto the surface (57). Prior to experiments, the microscope glass surfaces were sonicated in ultrapure water, rinsed with ethanol, and then dried with ultra-high purity nitrogen. The samples were then washed with ultrapure water and allowed to dry for one hour in a laminar flow hood (1300 Series A2, Thermo Scientific). Once dry, a stereoscope was used to locate and assess fibril deposition prior to AFM experiments.

Amplitude modulation (AM-) imaging and fast force mapping (FFM-) experiments were performed on a Cypher ES AFM (Asylum Research, Oxford Instruments). Silicon nitride cantilevers AC240 and Biolever-mini BL-AC40TS sourced from Oxford Instruments were used for AM imaging in air and liquid, respectively. These cantilevers have a nominal spring constant of kc = 2.0 Nm^-1^ and 0.1 Nm^-1^, and radii of r = 7 nm and 8 nm, respectively. Image and force measurements were acquired at 25° C. The glass microscope slide surface with deposited collagen fibrils was mounted on a steel puck within the AFM box (a sealed enclosure). Samples were first imaged in air to locate and characterize the collagen fibrils. Before liquid experiments, the spring constant of the cantilever was first determined in air using the GetReal function in the Asylum Research AFM software, which is a combination of the Sader (58) and thermal noise method. The inverse optical lever sensitivity (InVOLS) was then determined by averaging the results of ten force curves on a sapphire surface in PBS buffer. Liquid experiments were completed in a droplet of approximately 0.25 mL of fresh 10 mM PBS buffer, pH 7.4 solution on the glass microscope slide surface at 25° C. To minimize thermal drift, this setup was allowed to equilibrate for at least one hour in the AFM prior to imaging. To facilitate obtaining high quality AM images in liquid conditions, the tip was photothermally excited at its resonance frequency using blueDrive (Asylum Research, Oxford Instruments).

### Fast Force Mapping (FFM)

FFM experiments were performed immediately following AM-AFM imaging on the same fibrils characterized in air and liquid. The same BL-AC40TS AFM probe as referenced above was used for all nanoindentation experiments. Force maps were acquired at a scan size of 1 μm x 1 μm with a pixel resolution of 256 x 256 points, Z-rate of 20 Hz, indentation speed of 16 μm/sec, and a setpoint of 2 nN. This setpoint was used specifically to ensure that the indentation did not exceed 15% of the fibrils’ total height. The indentation moduli of the fibrils were then determined from the force maps produced using the analysis of the matrix of force curves as described below.

### Image Analysis

#### Periodicity

2D Fourier Transforms were performed on the phase channel of the AM images of collagen fibrils in both air and liquid using the Gwyddion software (59). Profiles were then drawn on the transformed images which then revealed the periodicity of the fibrils.

#### Overlap–Gap Step Height

For determination of the overlap–gap step height from the AM images, sine curves bearing the periodicity of the respective fibrils were fitted to height profiles along the apex of the corresponding fibril. The maxima/minima values of the sin curve were strictly used to determine the corresponding overlap/gap height values used along the fibrils. Adjacent overlap and gap values were then subtracted to obtain the step heights and this was performed across an entire height profile (Figure S4). Furthermore, some chondroitinase treated fibrils imaged in air showed signs of debris (aggregates from chondroitinase stock solution) occurring along the height profiles, these areas were not taking into consideration for determining the average height of the fibril or step height (Figure S5). When the treated fibrils were immersed in PBS, visual inspection revealed that there was no evidence that the debris remained on the fibrils.

### Post-Processing Analysis

FFM data sets were analyzed using in-house Matlab (R2018b) scripts. In brief, indentation moduli were calculated for each force curve iteratively using the methods detailed by Andriotis and coworkers (57). The resulting 256×256 matrix of FFM indentation moduli were then reconstructed in Cartesian space as a scaled colormap (Figure 3A–B). Box and whisker plots were derived from a subset of five FFM images for each treatment. Here, force curve data at the height apex along the fibril longitudinal axis was extracted (256 moduli) and pooled among treatments. For this study, the apex corresponds to the values along a straight line created between two of the highest values at opposite ends of the fibril. A one-way analysis of variance (ANOVA) was performed, and the moduli data from each treatment condition are shown as a box and whisker in the main text Figure 3C.

## Abbreviations

ECM: extracellular matrix;
PG: proteoglycan;
GAG: glycosaminoglycan;
α_2_I: I-domain of integrin α_2_ subunit;
CMP: collagen model peptide;
AFM: atomic force microscopy;
ELISA: enzyme-linked immunosorbent assay;
EDTA: ethylenediaminetetraacetic acid;
BSA: bovine serum albumin;
AM-AFM: amplitude-modulation AFM;
FFM-AFM: fast force mapping AFM;
DMMB: dimethylmethylene blue;
InVOLS: inverse optical lever sensitivity;
ANOVA: analysis of variance.

## Supporting Material

Figures S1-7 can be found in the Supporting Material: RothBaum_SI_ProteinScience.pdf.

## Acknowled gem ents

The authors acknowledge Dr. Joseph Freeman and Michael Pellegrini of Rutgers University Dept. of Biomedical Engineering for assistance with rat tail tendon procedures and Jordan Elliott in the Baum lab for assisting with integrin expression. This work was supported by NIH grants GM045302 and GM136431 to JB and an Exploratory Research Seed Grant from Rutgers University to RH.

## Notes

### Competing Interest Statement

The authors have declared no competing interest.

