## Supplementary Information for "Probing the effect of glycosaminoglycan depletion on integrin interactions with collagen I fibrils in the native ECM environment"

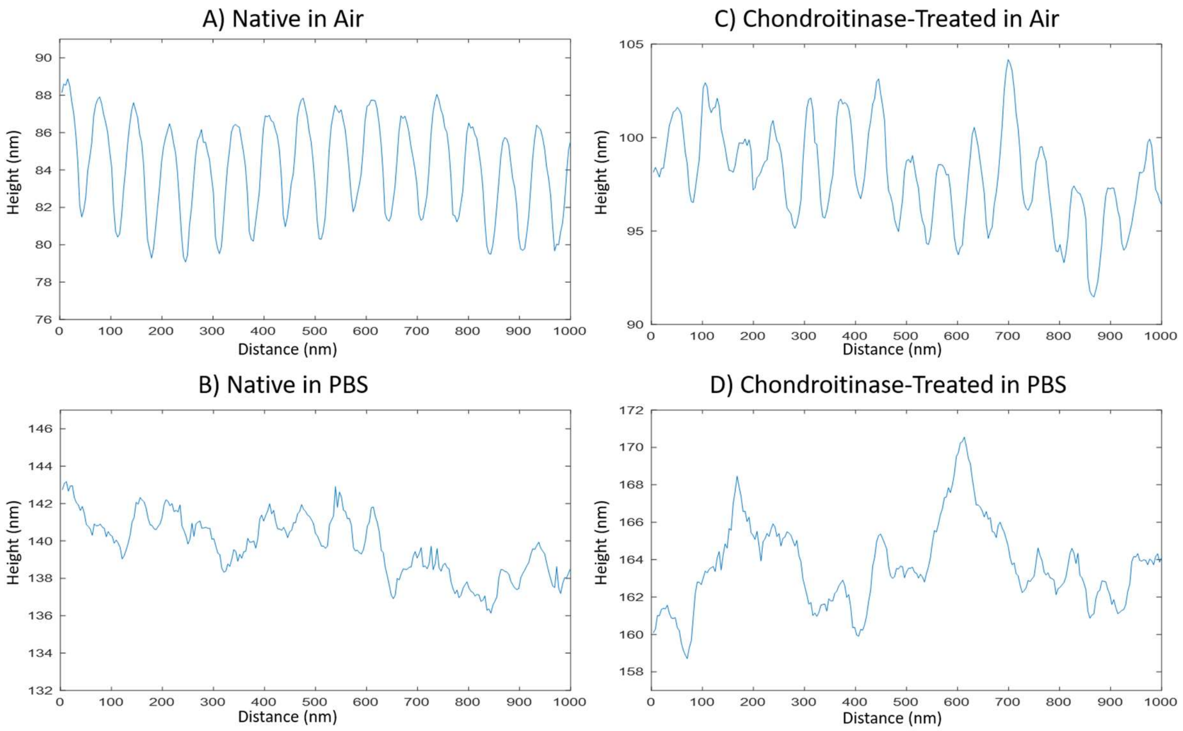


**Figure S1.** Full apex height traces from AM images along representative regions of (A, B) native and (C, D) chondroitinase-treated rat tail collagen fibrils in (A, C) air and (B, D) PBS, pH 7.4. In air, the fibrils have a clear repeating structure with distinct overlap and gap regions while in liquid this clarity is diminished and overlap and gap distinctions cannot be made. Visually there are still regions that resemble distinct overlap and gap regions albeit not throughout the entire profile.

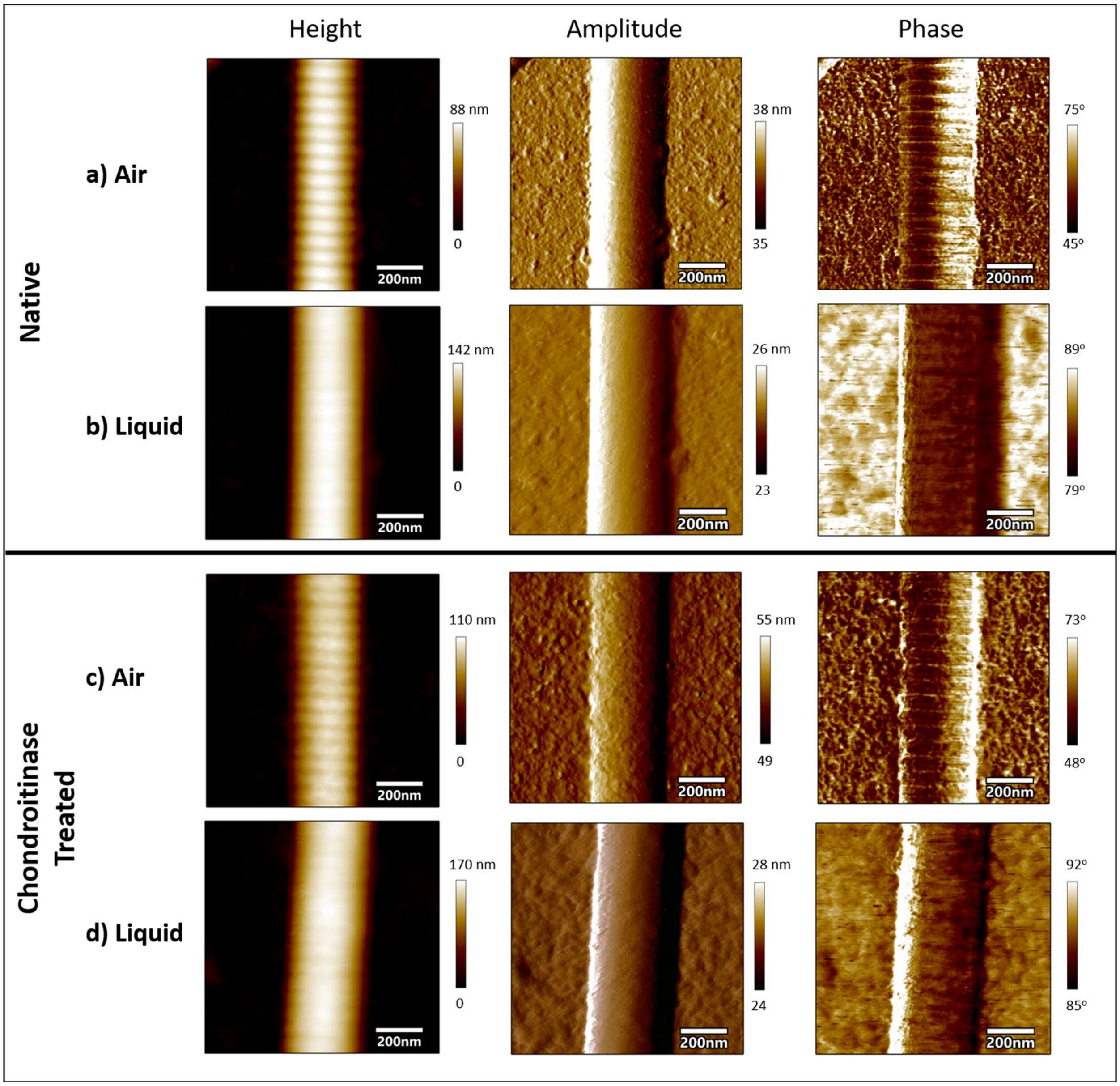


**Figure S2.** Uncropped AM images of native (A,B) and chondroitinase-treated (C,D) collagen fibrils in both air (A,C) and liquid (B,D) conditions. The height images are the same data as presented in Figure 2 in the main text. The corresponding amplitude and phase channels are displayed for each fibril in each condition.

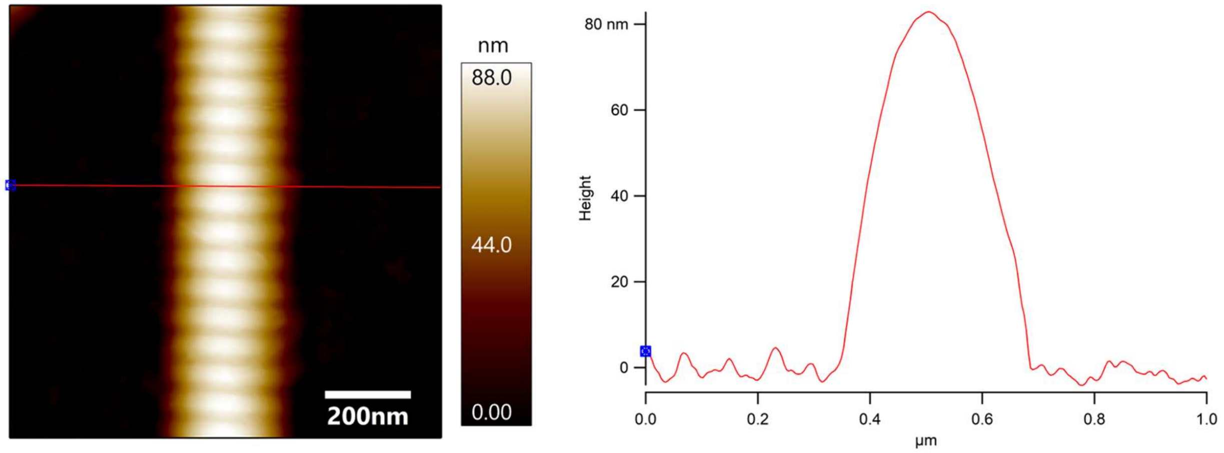


**Figure S3**. Height profile of a native collagen fibril in air. Heights were calculated by averaging the height values along the apex of the fibril. The calculated height of this fibril is 82.1 nm.

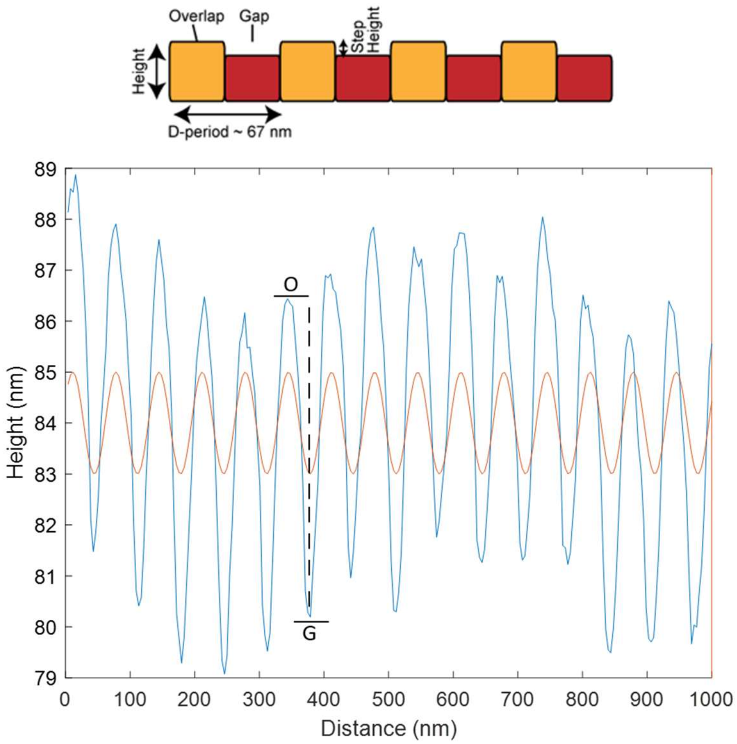


**Figure S4**. Cartoon schematic of the alternating heights between overlap and gap regions along the collagen fibril and a height profile along the apex of a native collagen fibril in air (blue curve) along with a sine curve fitted to the periodicity of the fibril (orange). In order to determine the overlap-gap step height, the height value corresponding to the maxima of the sin curve is considered the overlap height (letter “O”) is subtracted by the neighboring height value corresponding to the minima of the sine curve which is considered to be the gap height (letter “G”). This is repeated across the entire fibril and performed for all fibrils in air.

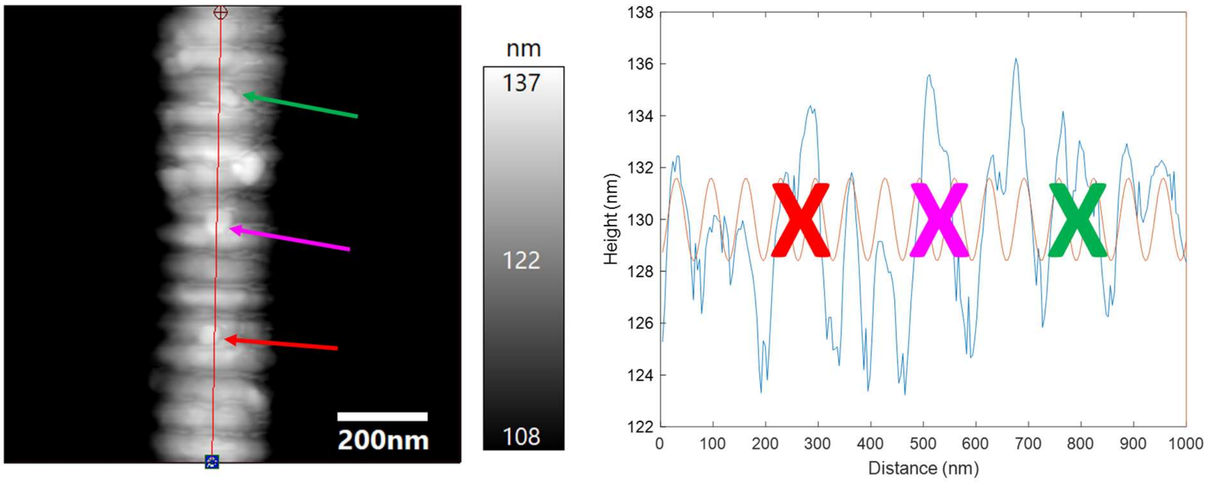


**Figure S5**. Height image of a chondroitinase treated fibril in air (left) and its height profile fitted with a sin curve similar to Figure S4 (right). In the image, there are signs of debris on the fibril as a result of the chondroitinase treatment (arrows). The debris can be distinguished on the fibril image as well as on the corresponding height profile. Since the debris disallows for the distinction of certain overall and gap regions and does not represent the true topography of the fibril these area's (marked by "x") were not accounted for step-height.

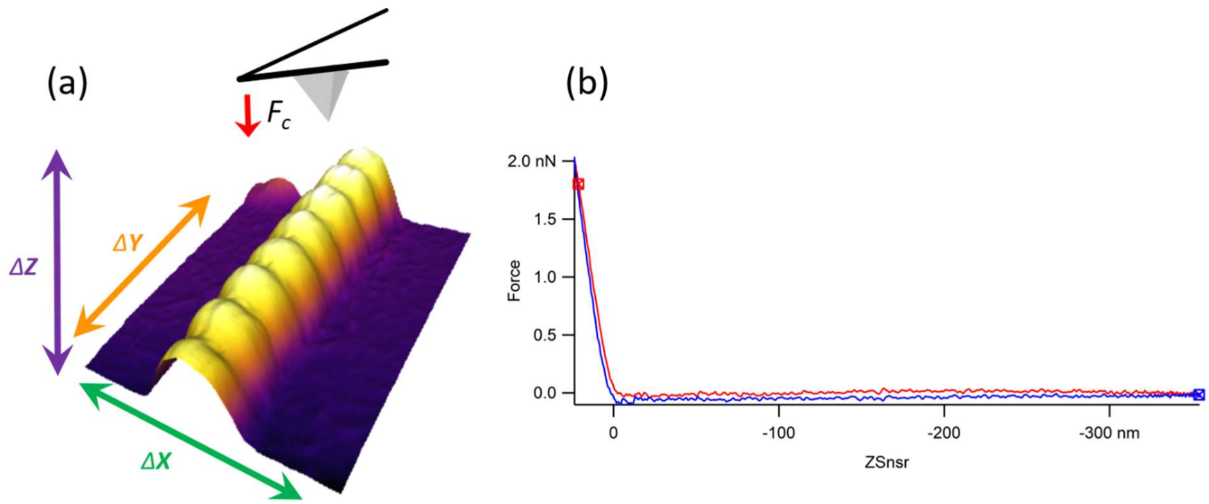


**Figure S6.** Fast Force Mapping. (A) Once a fibril is imaged and visualized in standard tapping mode the same exact area is divided into a 256 x 256 pixel grid, and a force curve is performed at each pixel. A representative force curve from a native fibril in liquid is shown in (B). The contact force for each coordinate is extracted and local modulus calculated. These moduli are then replotted in space to yield force maps such as that presented in Figure 3 of the main text.

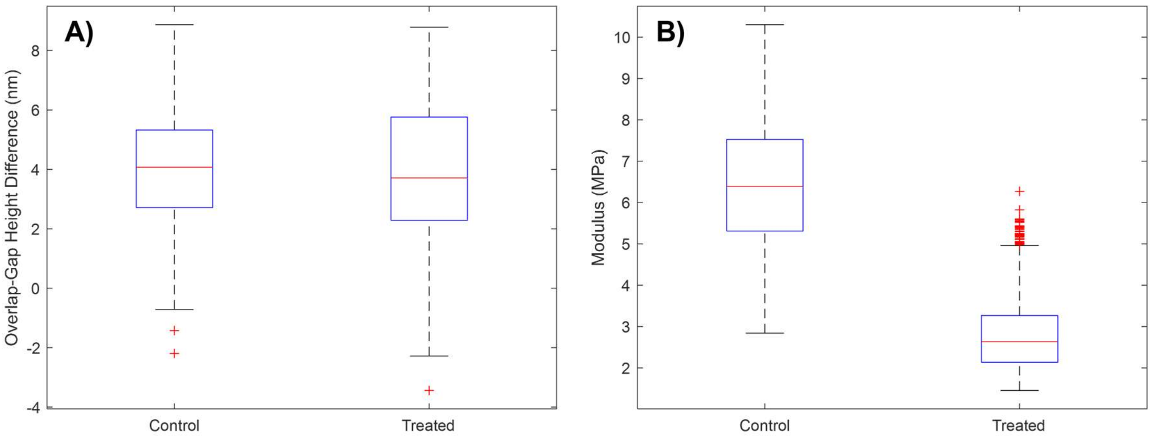


**Figure S7.** Boxplots showing the distribution of overlap-gap step height differences (A), and modulus values (B) between the control and chondroitinase treated fibrils. The step heights are not significantly different indicating no structural differences while the modulus values are significantly different. Red symbols indicate outliers of the boxplot.
